# Digital twinning of cardiac electrophysiology for congenital heart disease

**DOI:** 10.1101/2023.11.27.568942

**Authors:** Matteo Salvador, Fanwei Kong, Mathias Peirlinck, David W. Parker, Henry Chubb, Anne M. Dubin, Alison Lesley Marsden

## Abstract

In recent years, blending mechanistic knowledge with machine learning has had a major impact in digital healthcare. In this work, we introduce a computational pipeline to build certified digital replicas of cardiac electrophysiology in pediatric patients with congenital heart disease. We construct the patient-specific geometry by means of semi-automatic segmentation and meshing tools. We generate a dataset of electrophysiology simulations covering cell-to-organ level model parameters and utilizing rigorous mathematical models based on differential equations. We previously proposed Branched Latent Neural Maps (BLNMs) as an accurate and efficient means to recapitulate complex physical processes in a neural network. Here, we employ BLNMs to encode the parametrized temporal dynamics of in silico 12-lead electrocardiograms (ECGs). BLNMs act as a geometry-specific surrogate model of cardiac function for fast and robust parameter estimation to match clinical ECGs in pediatric patients. Identifiability and trustworthiness of calibrated model parameters are assessed by sensitivity analysis and uncertainty quantification.

## 1 Introduction

The combination of physics-based and statistical modeling in cardiovascular medicine has the potential to shape the future of cardiology [9]. In this framework, a synergistic use of multiphysics and multiscale mathematical models for cardiac function [13, 18, 39, 41, 57] and machine learningbased methods, such as Gaussian processes emulators [30, 37, 59] and Neural Networks (NNs) [35, 47], enables the design of efficient computational tools that are compatible with the computer resources and time frames required in clinical applications. In the foreseeable future, a continuous, bi-directional interaction between patient-specific data and Artificial Intelligence-enriched computer models incorporating biophysically detailed and anatomically accurate knowledge would enable the vision of precision medicine [34, 38, 44]. Personalized treatment and surgical planning may be delivered by leveraging different mathematical methods, such as sensitivity analysis, parameter inference and uncertainty quantification [19, 25, 49, 54].

Several mathematical tools have been proposed to better understand and treat different groups of adult cardiac pathologies [34]. Electrophysiology simulations play an important role in the assessment of rhythm disorders. They are used for cardiac resynchronization therapy [58], arrhythmia risk stratification [2, 50], and definition of optimal ablation strategies [43]. Nevertheless, in silico numerical simulations and treatment modalities in pediatrics and congenital heart disease are less common or even not established [8, 24, 62].

Congenital heart defects (CHDs) are the most common birth defects and are characterized by cardiac anatomical abnormalities that can severely impact cardiac function [29]. Patients with CHDs often have a unique and peculiar combination of cardiac defects that warrant personalized treatment planning in clinically-relevant time frames. Digital twinning of cardiac function thus holds particular promise for these patients [62].

In this work we introduce a novel digital twin of a pediatric patient with hypoplastic left heart syndrome (HLHS), a complex form of CHD where the left ventricle of the patient is severely underdeveloped, leading to a number of morbidities and elevated mortality risk. Our pipeline seamlessly combines:

- Semi-automatic segmentation and mesh generation tools suited for pediatric patients with CHD [27]
- A physiologically-based mathematical formulation of cardiac electrophysiology deriving from the monodomain equation [44, 51] coupled with the ten Tusscher-Panfilov [63] ionic model
- A recently proposed scientific machine learning method, namely Branched Latent Neural Maps (BLNMs) [53], to build an accurate and efficient dynamic surrogate model of cardiac function
- Shapley effects [56] and Hamiltonian Monte Carlo (HMC) [5, 21] to perform patient-specific sensitivity analysis, fast and robust parameter estimation with uncertainty quantification while matching clinical 12-lead electrocardiograms (ECGs)

This digital twin can be employed to simulate different scenarios of clinical interest in silico, as HLHS patients may experience different forms of electrophysiological comorbidities [14], including ventricular arrhythmias and dyssynchrony [62]. Therefore, personalized electrophysiology simulations may provide virtual preand post-operative guidance in this understudied patient cohort [32].

## 2 Results

In Figure 1 we depict our computational pipeline to build digital twins of cardiac electrophysiology for congenital heart disease patients. This pipeline covers all the relevant aspects of digital twinning: image segmentation and mesh generation, mathematical and numerical physics-based modeling, surrogate model training, sensitivity analysis and robust parameter calibration with uncertainty quantification.

**Figure 1:**
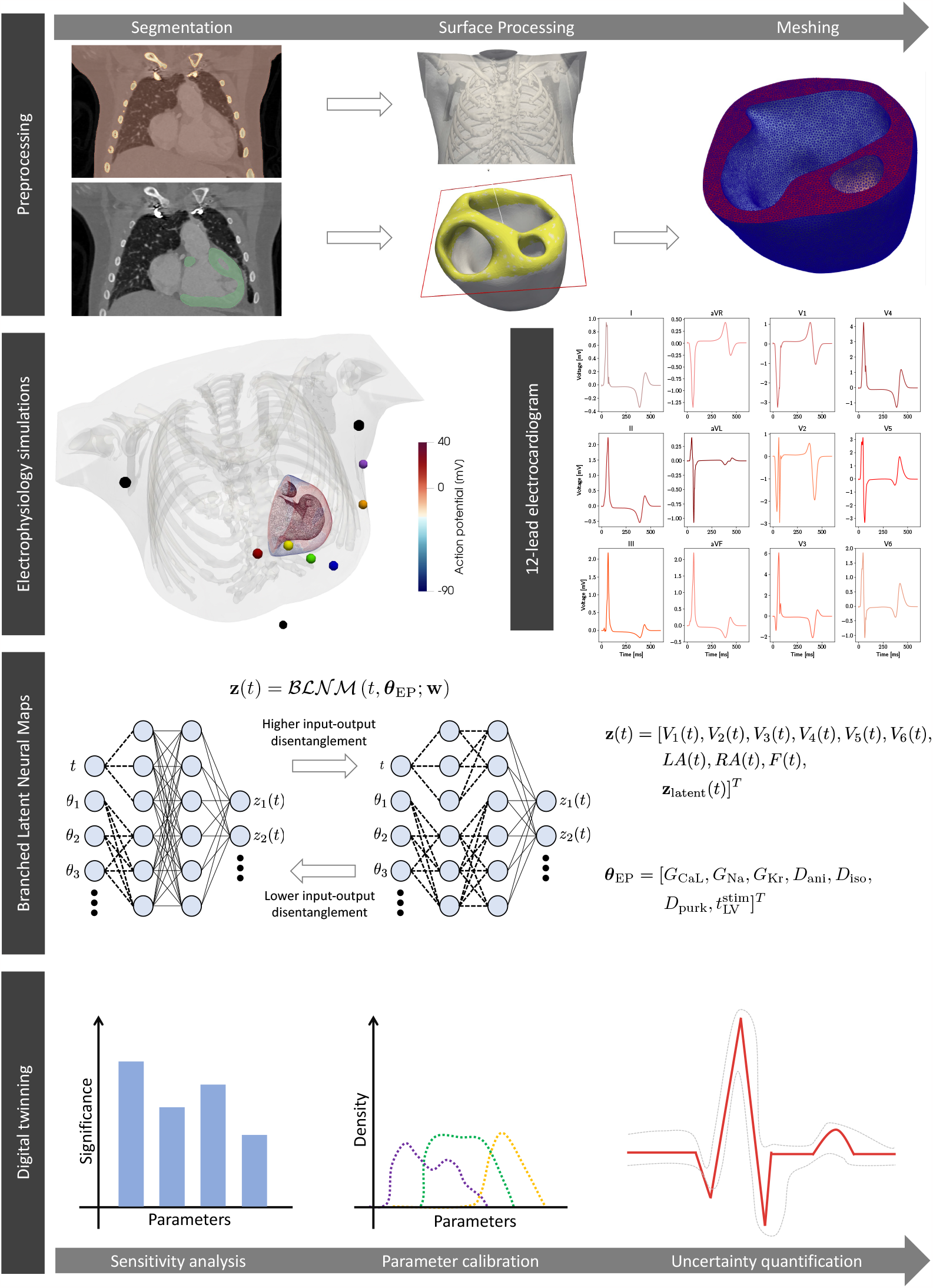
Sketch of the computational pipeline. We reconstruct the patient-specific geometry with HLHS from imaging. We generate a dataset of electrophysiology simulations encompassing cell-to-organ variability in model parameters. We train a BLNM that effectively reproduces 12-lead ECGs while covering model variability. We employ the BLNM for digital twinning.

### 2.1 Pre-processing

Figure 1 (first row) shows the heart-torso model of a 7-year-old female pediatric patient with HLHS constructed from the computerized tomography (CT) scan of the patient using our semi-automatic model construction pipeline [27].

Electrophysiology simulations

### 2.2 Cardiac electrophysiology modeling

We run 200 numerical simulations on the patient-specific heart-torso geometry (see Figure 1, second row), spanning seven relevant electrophysiology parameters of the physics-based model at the microscopic scale and organ level. We collect the corresponding in silico 12-lead ECGs. In Table 1 we report descriptions, ranges, and units for the seven model parameters that we explore via latin hypercube sampling for the dataset generation.

**Table 1:** Parameter space for cardiac electrophysiology sampled via latin hypercube for the numerical simulations performed with the physics-based mathematical model.

| Parameter | Description | Range | Units |
| --- | --- | --- | --- |
| $G_{CaL}$ | Maximal $Ca^{2+}$ current conductance | [1.99e-5, 7.96e-5] | cm ms<sup>-1</sup> $\mu F^{-1}$ |
| $G_{Na}$ | Maximal $Na^+$ current conductance | [7.42, 29.68] | nS pF<sup>-1</sup> |
| $G_{Kr}$ | Maximal rapid delayed rectifier current conductance | [0.08, 0.31] | nS pF<sup>-1</sup> |
| $D_{ani}$ | Anisotropic conductivity | [0.008298, 0.033192] | mm<sup>2</sup> ms<sup>-1</sup> |
| $D_{iso}$ | Isotropic conductivity | [0.002766, 0.011064] | mm<sup>2</sup> ms<sup>-1</sup> |
| $D_{purk}$ | Purkinje conductivity | [1.0, 3.5] | mm<sup>2</sup> ms<sup>-1</sup> |
| $t_{LV}^{stim}$ | Purkinje left bundle stimulation time | [0, 100] | ms |

In Figure 2 we depict the ensemble of the resulting in silico 12-lead ECGs together with the clinical recordings. We point out that the patient-specific 12-lead ECGs are contained within the pseudopotentials variability spanned by the electrophysiology simulations, manifesting various morphologies in the QRS complex, that is ventricular depolarization, and T wave, that is ventricular repolarization. The patient diagnosis reports rhythm disorders, atrial enlargement, left and right ventricular hypertrophy, along with severe abnormalities in the ECGs. Specifically, there are signs of prolonged PR interval, ST segment depression and T wave inversion.

**Figure 2:**
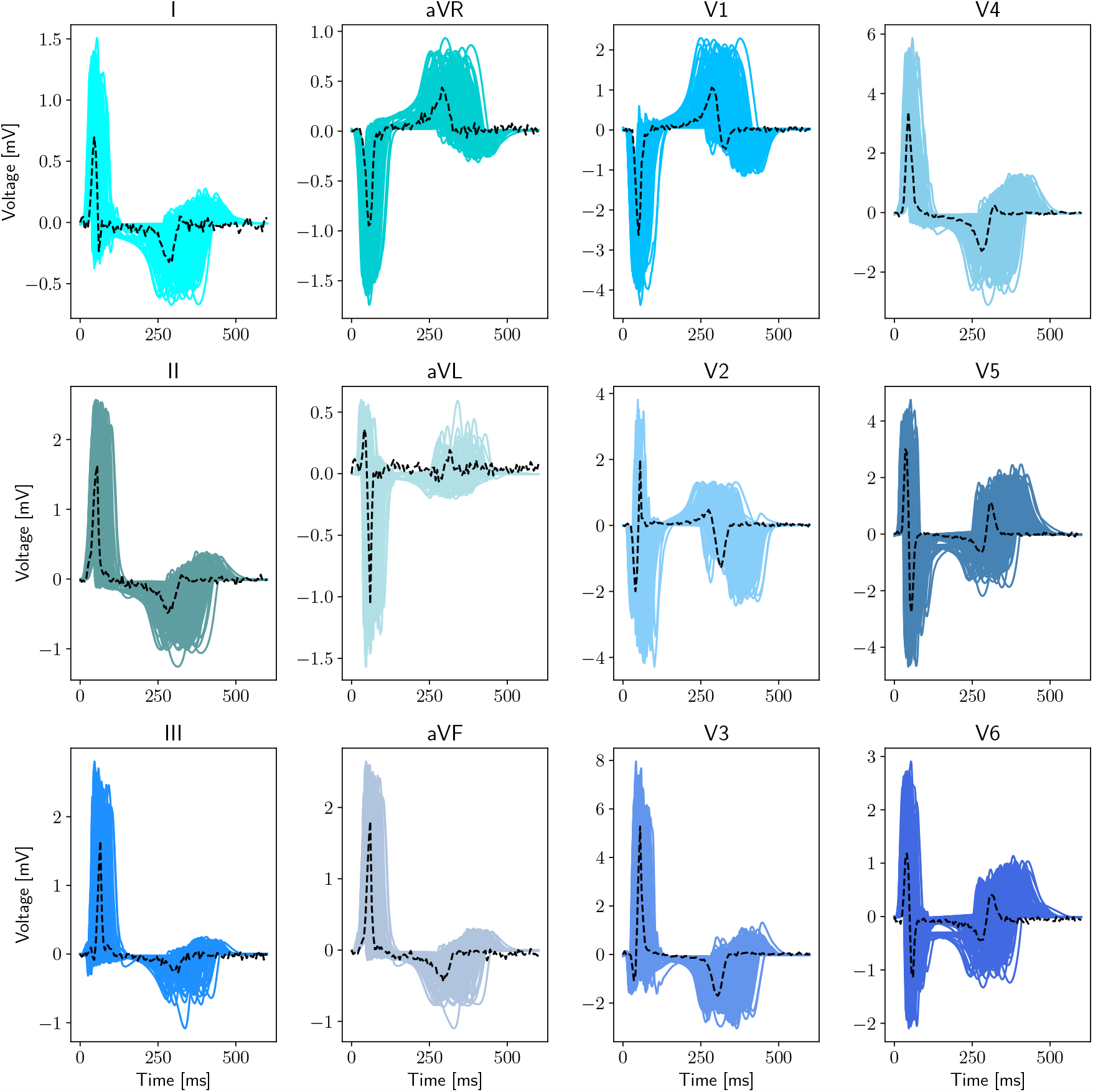
Physics-based electrophysiological modeling dataset generation. Full dataset containing 200 in silico precordial and limb leads recordings (blue, solid) and patient-specific 12-lead ECGs (black, dashed).

In Figure 3 we show the simulated spatio-temporal transmembrane potential evolution on the patient-specific pediatric model for a single instance of model parameters. Specifically, we always simulate the sinus rhythm behavior over a cardiac cycle. Figure 3 focuses on the ventricular depolarization phase, where the electric signal propagates from the 1D Purkinje network at the two endocardia towards the myocardium, as well as the ventricular repolarization phase, when the transmembrane potential comes back to its resting state (i.e. approximately -90 mV).

**Figure 3:**
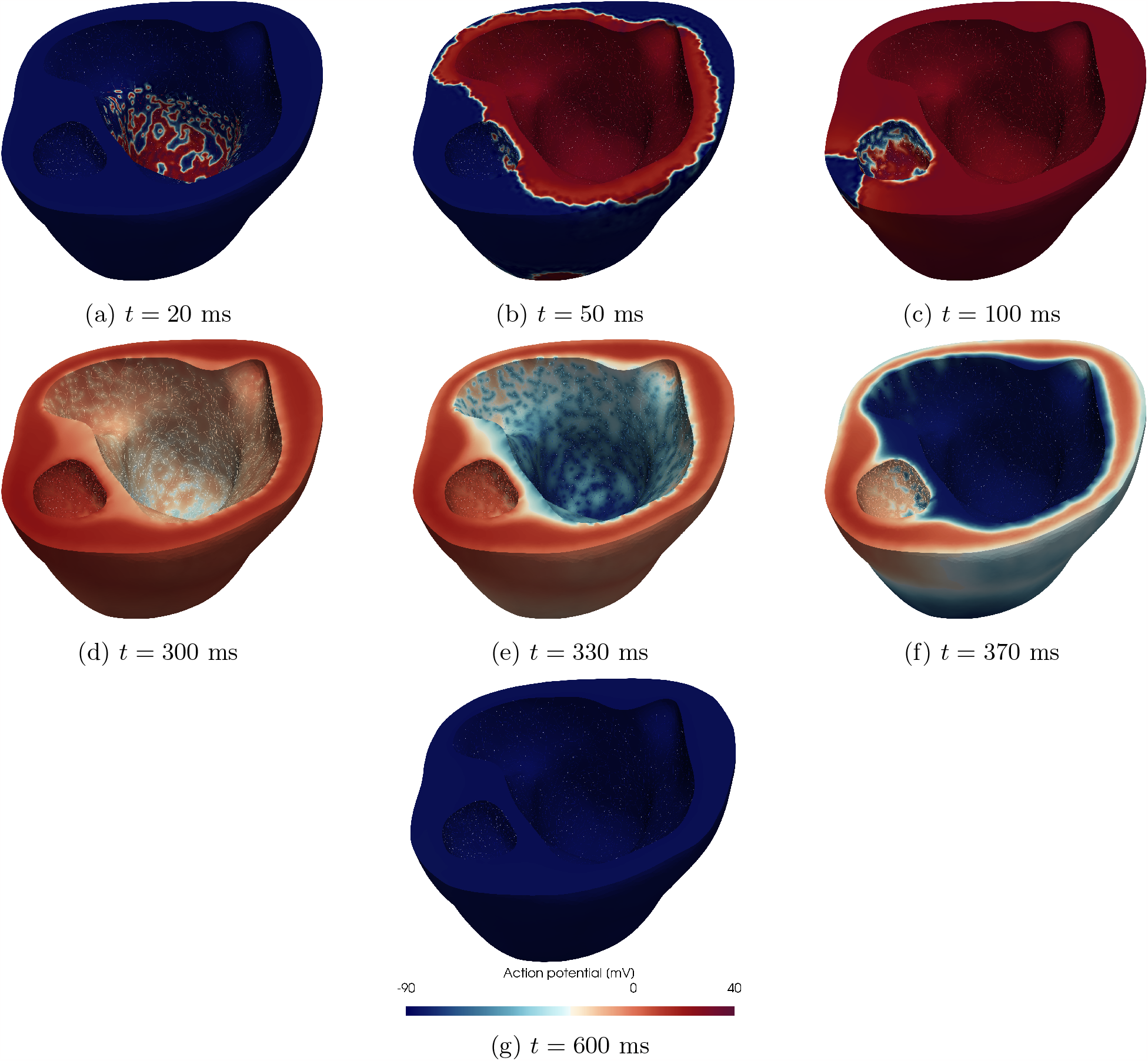
Physics-based electrophysiological modeling. Spatio-temporal membrane action potential evolution for one electrophysiology simulation in the dataset performed on the HLHS pediatric patient.

### 2.3 Branched Latent Neural Maps

We train BLNMs, which are represented by feedforward partially-connected NNs, to encode the temporal dynamics of the 12-lead pseudo-ECGs computed with the physics-based mathematical model while also covering model variability from the cellular to the tissue level (see Figure 1, third row). Once trained, BLNMs act as a surrogate model for cardiac electrophysiology function that can be queried on new parameter instances to provide faster than real-time in silico 12-lead ECGs. In order to identify the optimal set of BLNM hyperparameters, which are the number of layers, number of neurons, number of states, and disentanglement level in the NN structure, we employ a *K*fold (*K* = 5) cross validation over 150 multiscale physics-based electrophysiology simulations. The hyperparameter search space is given by a four-dimensional hypercube, where we run 50 instances of Latin Hypercube Sampling and we pick the BLNM configuration providing the lowest generalization error. For each configuration of hyperparameters, we sample the dataset with a fixed time step of Δ*t* = 5 ms and we perform 10,000 iterations of the second-order Broyden-Fletcher-Goldfarb-Shanno (BFGS) optimizer. In Table 2 we report the initial hyperparameters ranges for tuning and the final optimized values.

**Table 2:** Branched Latent Neural Map hyperparameter tuning. Original hyperparameter ranges and optimized hyperparameter values for the final training stage.

| BLNM |  |  | Hyperparameters |  | Trainable parameters |
| --- | --- | --- | --- | --- | --- |
|  | layers | neurons | number of states | disentanglement level | # parameters |
| tuning | {1 ... 8} | {10 ... 30} | {9 ... 12} | {1 ... $N_{\text{layers}}$} | |
| final | 7 | 19 | 10 | 2 | 2,398 |

Then, we train a final optimized BLNM encompassing the whole dataset of 150 multiscale physics-based electrophysiology simulations using 50,000 BFGS iterations. The non-dimensional Mean Square Error (MSE) on a testing set comprised of 50 additional numerical simulations unseen during the training stage, and Latin Hypercube sampled from the same parameter space in Table 1, is equal to 5 · 10^*−*4^.

### 2.4 Parameter estimation

We employ the optimized BLNM to find an initial guess for the seven model parameters that results in a computational pseudo-ECG that best matches the clinically observed 12-lead ECG dynamics of the CHD patient. The estimated model parameters are reported in Table 3.

**Table 3:** Parameter estimation. Calibration with the optimized BLNM for cell-to-organ level model parameters of the physics-based mathematical model. The MSE between the BLNMs predictions and the clinical recordings, in non-dimensional form, is 0.16.

| Parameter | Value | Units |
| --- | --- | --- |
| $G_{\text{CaL}}$ | 2.94e-5 | $\text{cm ms}^{-1} \mu\text{F}^{-1}$ |
| $G_{\text{Na}}$ | 15.58 | $\text{nS pF}^{-1}$ |
| $G_{\text{Kr}}$ | 0.15 | $\text{nS pF}^{-1}$ |
| $D_{\text{ani}}$ | 0.03 | $\text{mm}^2 \text{ms}^{-1}$ |
| $D_{\text{iso}}$ | 0.01 | $\text{mm}^2 \text{ms}^{-1}$ |
| $D_{\text{purk}}$ | 1.96 | $\text{mm}^2 \text{ms}^{-1}$ |
| $t_{\text{LV}}^{\text{stim}}$ | 43.3 | ms |

Even though the relative heart orientation and lead placements significantly influence ECGs [66], and as such may require additional parameters to calibrate, the information retrieved from the CT scan and patient diagnosis allow us to determine these quantities with a small degree of uncertainty. This motivates our focus in the estimation process on the cell-to-organ level electrophysiology model parameters which are assessed as important in previous studies [10, 60].

### 2.5 Sensitivity analysis

Starting from the parameter calibration shown in Table 3, we compute Shapley effects for each model parameter for cardiac electrophysiology, assuming independence among them as they act in different terms and equations of the physics-based mathematical model (see Figure 1, third row). In Figure 4 we show how each parameter contributes in matching electrophysiology simulations with the clinical 12-lead ECGs, i.e. in the minimization of the MSE between BLNM outputs and our patient-specific observations. The sodium current conductance *G*_Na_ plays a dominant role, followed by the L-type calcium ion channel conductance *G*_CaL_ and the different conductivities *D*_ani_, *D*_iso_ and *D*_purk_. Noteworthy, the interventricular activation dyssynchrony 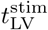 plays a minor role in the calibration process. This is motivated by the dimension of the right ventricle, which mostly dictates the activation sequence with respect to the small (underdeveloped) left ventricle.

**Figure 4:**
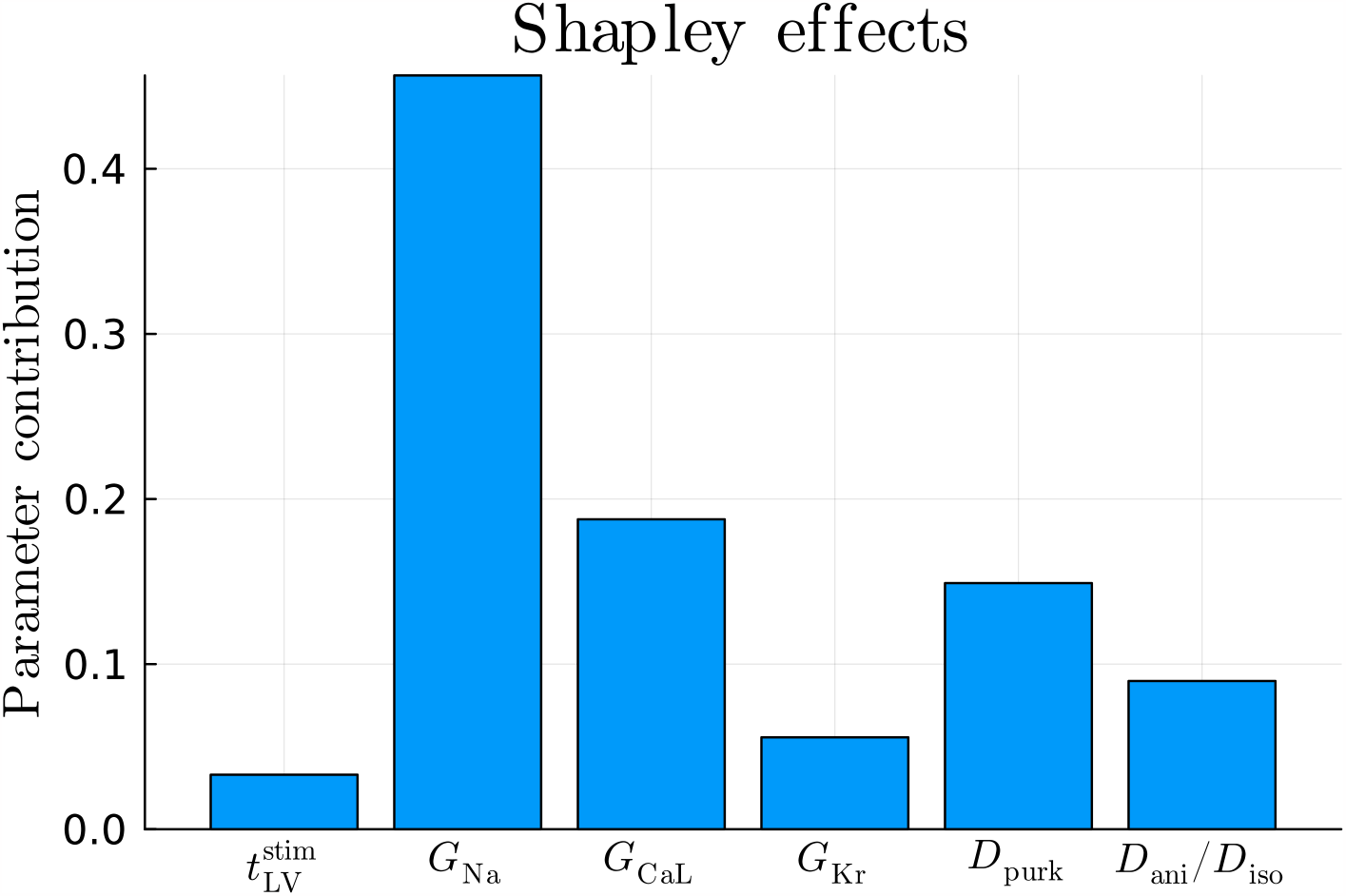
Sensitivity analysis for the seven model parameters encoded in the BLNM via Shapley effects.

### 2.6 Uncertainty quantification

In Figures 5 and 6 we show the results of our inverse uncertainty quantification, where we quantify how uncertainty in matching 12-lead ECGs propagates to uncertainty in the estimated model parameters. We account for both BLNM surrogate modeling error, via Gaussian processes (GPs), and measurement error in the clinical recordings.

**Figure 5:**
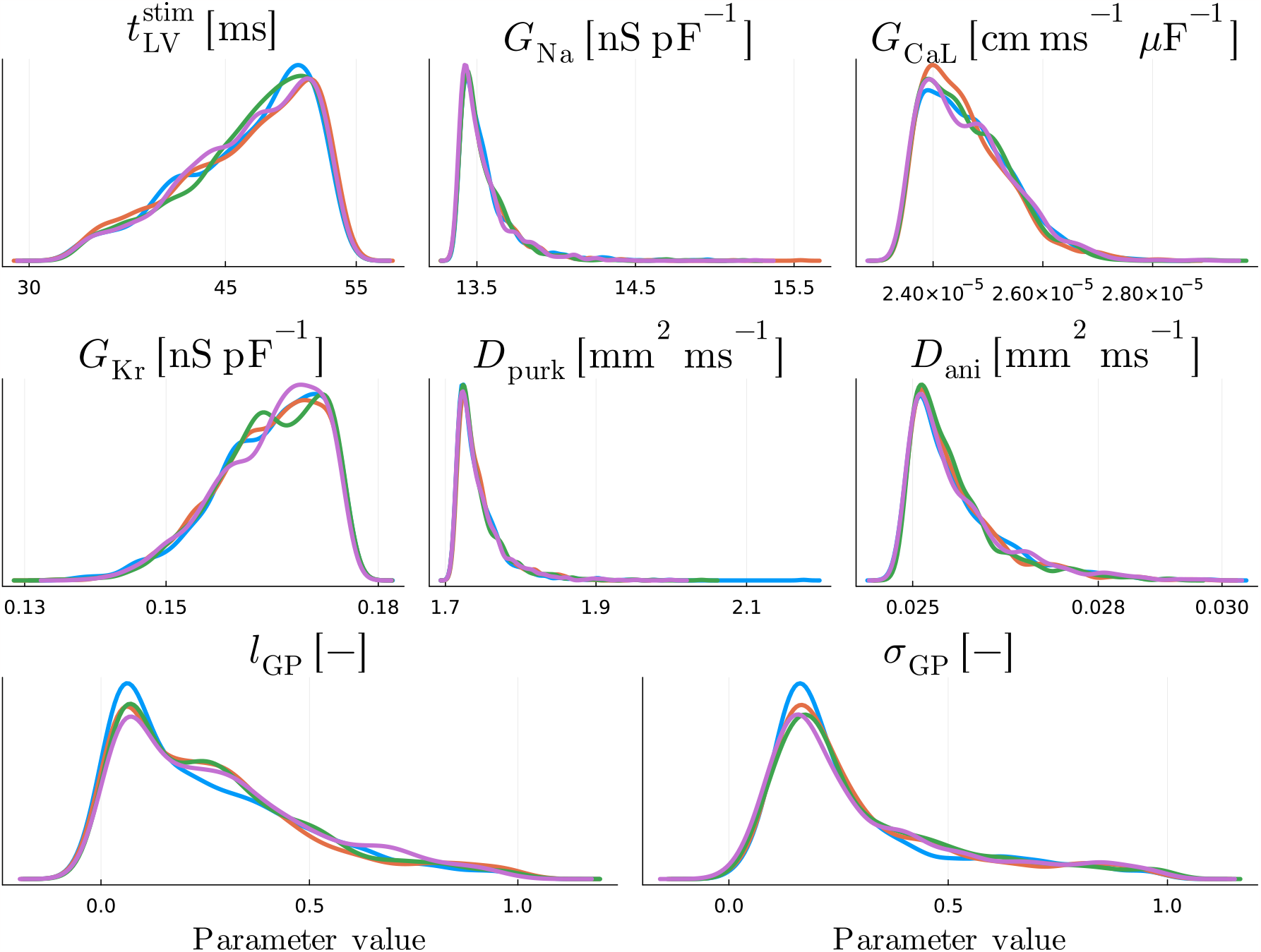
Inverse uncertainty quantification: parameter uncertainty. One-dimensional views of the posterior distribution. Different colors represent different HMC chains.

**Figure 6:**
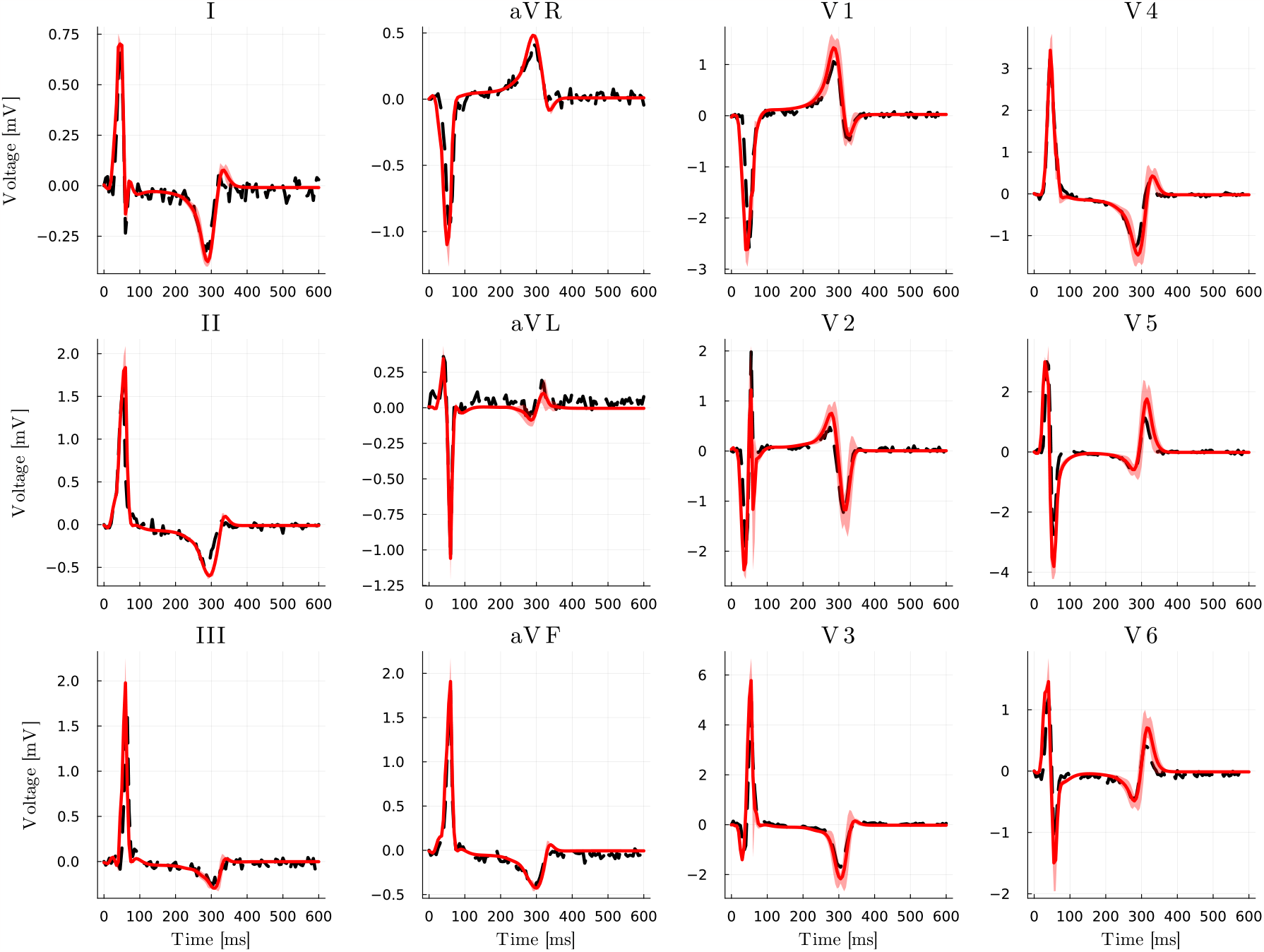
Inverse uncertainty quantification: matching clinical data. Clinical recordings (dashed, black) and mean estimation (red, solid) of 12-lead ECGs for the HLHS pediatric patient via HMC. Light red encompasses the variability between mean minus/plus five standard deviations.

From Figure 5, we see that the posterior distributions of all model parameters ***θ***_EP_, along with the correlation length *l*_GP_ and amplitude *σ*_GP_, converged well towards highly similar unimodal distributions across all chains. The average value of 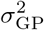 is approximately equal to 0.08, which is two orders of magnitude higher than the BLNM testing error (5·10^*−*4^), as the GP encodes the maximum BLNM uncertainty from each single lead individually and by also considering all possible correlations among the 12 leads, given the full covariance matrix in the multivariate normal distribution (see Section 3.6).

In Figure 6 we depict the clinical vs. in silico 12-lead ECGs, generated with the BLNM over the posterior distribution of model parameters. We see that the numerical simulations are in good agreement with the patient-specific recordings and show small variability between the five standard deviations from the average value.

### 2.7 In silico clinical trial

In Figure 7 we show the results of three numerical simulations in which we analyze different scenarios of clinical interest pre-operatively on the HLHS pediatric patient. Specifically, we depict activation and repolarization times for the electrophysiology simulation with the calibrated model parameters ***θ***_EP_ and by inducing either a left or a right bundle branch block, where the left (respectively, right) Purkinje fascicles are inhibited. We notice that, for this patient-specific case, the role of the Purkinje network in the left ventricle is very limited and that the activation sequence is highly similar with and without a full left bundle branch block.

**Figure 7:**
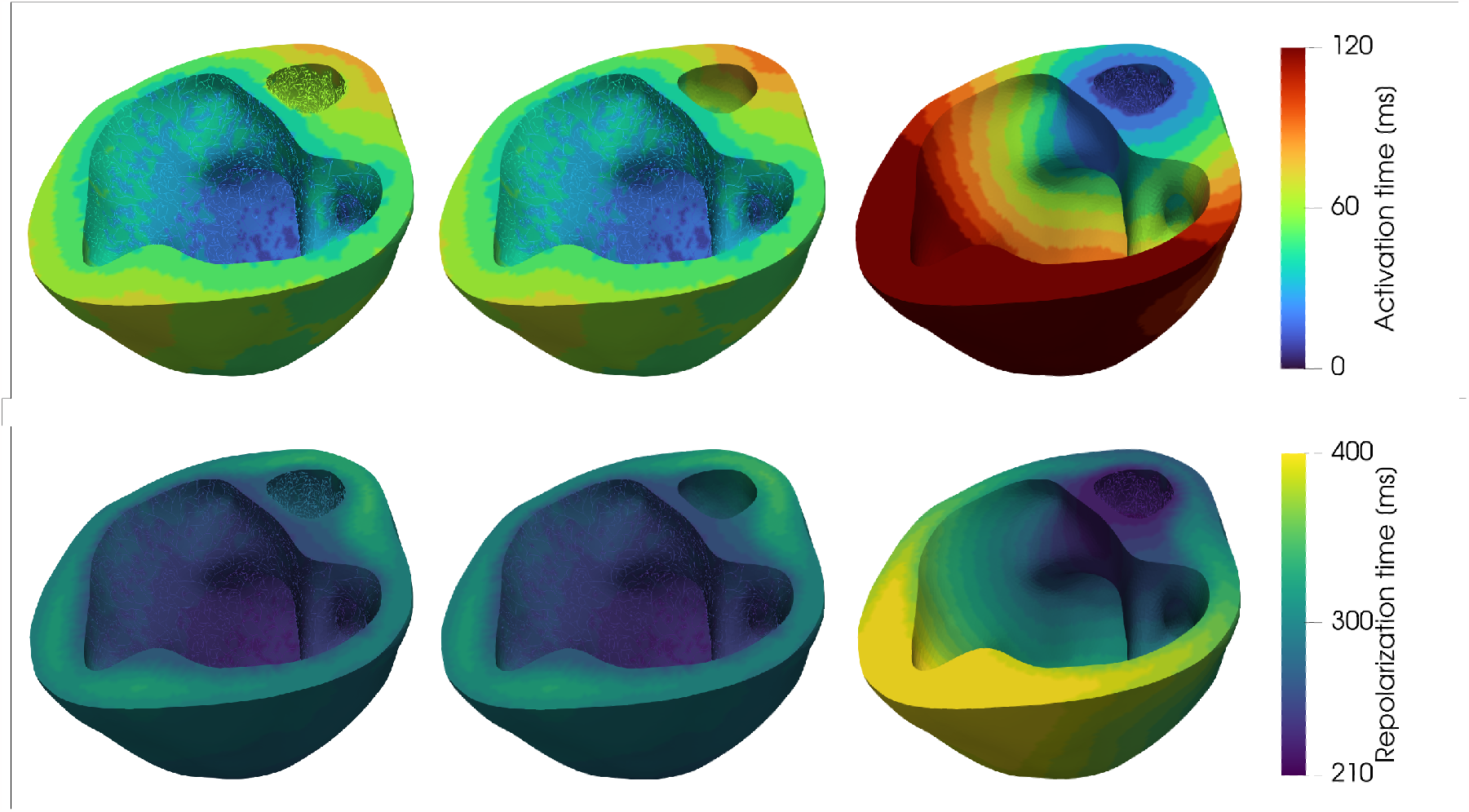
Running an in silico clinical trial. Activation (top) and repolarization (bottom) maps with personalized model parameters (left), left bundle branch block (center) and right bundle branch block (right).

### 2.8 Computational costs

In Table 4 we detail the computational costs and resources required by each step of the digital twinning process. The most expensive part resides in the physics-based computational electrophysiology modeling dataset generation, which makes use of high-performance computing given the stiffness and complexity of the underlying mathematical model. On the other hand, training a BLNM and employing it for robust Bayesian parameter estimation and sensitivity analysis are more tractable tasks that can be carried out within a few hours or minutes on a local machine. Using the physics-based mathematical model throughout parameter calibration with uncertainty quantification and sensitivity analysis would be computationally intractable and unaffordable given the extensive number of queries that Shapley effects and HMC require to show robustness and convergence in the provided results (see Methods section).

**Table 4:**
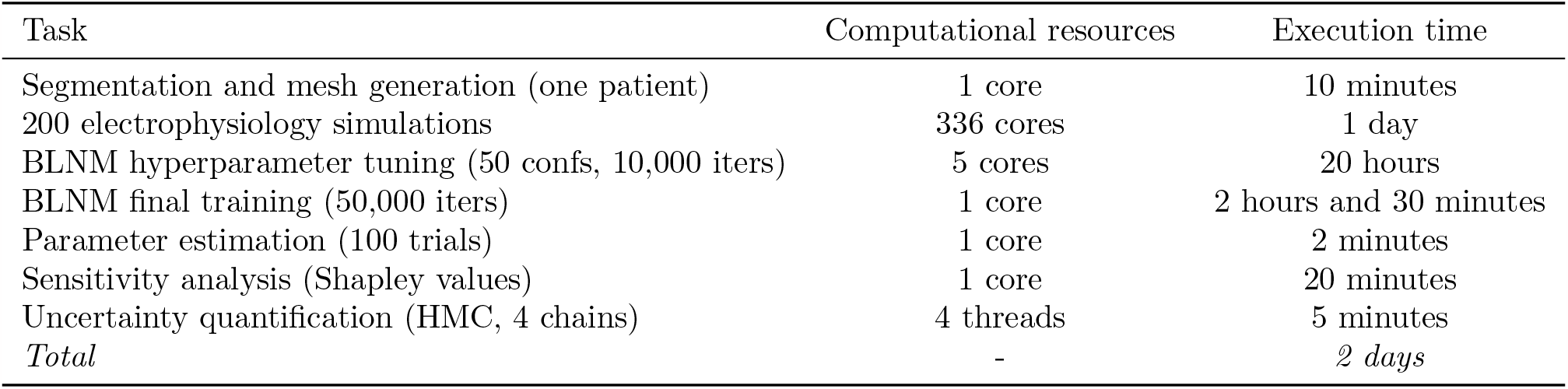
Computational resources. Summary of the computational times and resources to generate the electrophysiology simulations with the physics-based model, to train the BLNM, to compute Shapley values for sensitivity analysis and to perform Bayesian parameter estimation with uncertainty quantification on 12-lead ECGs.

## 3 Methods

### 3.1 Pre-processing

We reconstruct the heart-torso model from the CT scan of a 7-year-old female pediatric patient with HLHS in a semi-automatic manner [27]. Namely, we train a NN based on the classic UNet architecture [23] to automatically segment the myocardium from CT images. The UNet is trained using a publicly available dataset [67] that provided CT images and ground truth segmentation for 110 patients with age between 1 month and 40 years, combined with our private HLHS dataset containing the images and segmentation of 5 patients. Given the intrinsic segmentation challenges of cardiac structures in both young and CHD patients [40], we subsequently examine and improve the UNet-produced segmentations to more closely match with the CT scan. We automatically extract the surface meshes from the segmentations using the marching cube algorithm [31] and truncate the base myocardium above a manually identified plane to create a biventricular surface mesh. We subsequently use TetGen [55] to create the tetrahedral volumetric mesh with a maximum edge size of 1 mm [53, 62]. The torso model is created semi-automatically from the images using threshold- and region-growing-based segmentation methods. Images and associated clinical data were obtained under an IRB-approved protocol at Stanford University.

### 3.2 Cardiac electrophysiology modeling

We detail the physics-based mathematical model, along with its numerical discretization, that is employed to perform electrophysiology simulations on the HLHS pediatric patient.

#### 3.2.1 Mathematical model

We consider the monodomain equation [44] coupled with the ten Tusscher-Panfilov ionic model [63] to describe the electric behavior in the heart-Purkinje system. This system of differential equations reads:

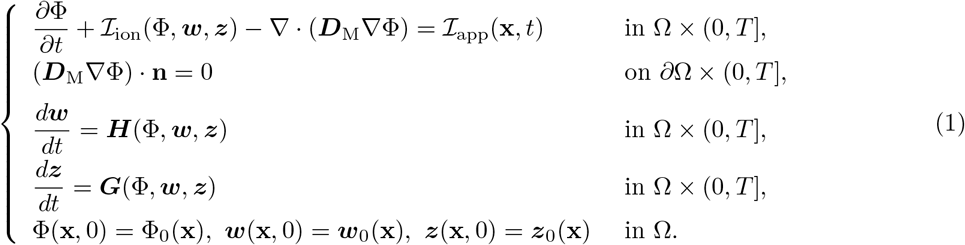

We always simulate a single heartbeat by fixing a final time *T* = *T*_HB_ = 600 ms. The computational domain Ω = Ω_purk_ Ω_myo_ is represented by the one-way coupled 1D Purkinje network and 3D biventricular patient-specific geometry. Transmembrane potential Φ defines the electric signal at the Purkinje and myocardial level. The ten Tusscher-Panfilov ionic model is endowed with 18 variables, which are split in two different subsets. First, there is a vector ***w*** = (*w*_1_, …, *w*_*M*_ ) (*M* = 12) of ion channel gating variables, which are probability density functions representing the fraction of open channels across the membrane of a single cardiac cell. Then, there is a vector ***z*** = (*z*_1_, …, *z*_*P*_ ) (*P* = 6) of concentration variables representing relevant ionic species, such as sodium *Na*^+^, intracellular calcium *Ca*^2+^ and potassium *K*^+^, which all play a major role in the metabolic processes [4], dictating heart rhythmicity or sarcomere contractility, and are generally targeted by pharmaceutical therapies. The specific mathematical formulation of the ten Tusscher-Panfilov ionic model defines the ordinary differential equations for ***H***(Φ, ***w, z***) and ***G***(Φ, ***w, z***), which describe the dynamics of gating variables and ionic concentrations respectively, along with the ionic current ℐ_ion_(Φ, ***w, z***) [63]. An external applied current ℐ_app_(**x**, *t*) fires the electric signal in the Purkinje fibers.

The diffusion tensor is expressed as ***D***_M_ = *D*_iso_**I** + *D*_ani_**f**_0_⊗**f**_0_ in Ω_myo_ and ***D***_M_ = *D*_purk_**I** in Ω_purk_, where **f**_0_ introduces the biventricular fiber field [42, 51]. *D*_ani_, *D*_iso_, *D*_purk_ ∈ ℝ^+^ dictate the anisotropic, isotropic and Purkinje conductivities, respectively.

The homogeneous Neumann boundary conditions prescribed at *∂*Ω define the condition of an electrically isolated domain, where **n** is the outward unit normal vector to the boundary.

Following [51], the extracellular potential Φ_e_ defining the ECGs is computed in each lead location **x**_e_ as:

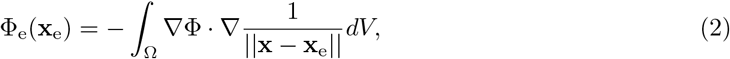

where *e* = {*V*_1_, *V*_2_, *V*_3_, *V*_4_, *V*_5_, *V*_6_ }and *e* = {*LA, RA, F}* define six precordial leads and three limb leads located on the pediatric patient-specific torso model, shown in colored and black dots in Figure 1 (second row), respectively. From these lead locations, we computationally reconstruct three bipolar limb leads as:

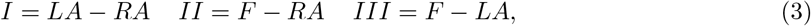

and three augmented limb leads as:

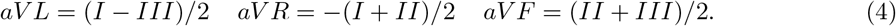

The resulting set *ECG* = {*V*_1_, *V*_2_, *V*_3_, *V*_4_, *V*_5_, *V*_6_, *I, II, III, aV L, aV R, aV F}* of computational pseudopotentials defines a comprehensive 12-lead ECG representation of the electrical activity in the patient-specific heart.

#### 3.2.2 Numerical discretization

We employ linear Finite Elements to discretize the spatial domain Ω in Equation (3.2.2). The tetrahedral tessellation defining the biventricular mesh has 933,916 cells and 158,277 DOFs, with a maximum mesh size of *h* = 1 mm. The left and right Purkinje bundles within the ventricular endocardia are generated by employing the fractal tree and projection algorithm proposed in [48], starting from the atrioventricular node. These left and right bundles are endowed with 14,820 elements (14,821 DOFs) and 67,456 elements (67,457 DOFs), respectively. Given the space resolution of the biventricular mesh, we apply non-Gaussian quadrature rules to recover convergent conduction velocities [62]. We consider a transmural variation of ionic conductances to differentiate epicardial, mid-myocardial and endocardial properties [63]. To solve Eq., we leverage an Implicit-Explicit time discretization scheme, where we first update the variables of the ionic model and then the transmembrane potential [13]. Specifically, in the monodomain equation, the diffusion term is treated implicitly and the ionic term is treated explicitly. The latter is discretized by means of the Ionic Current Interpolation scheme [28]. We prescribe the fiber distribution according to a Laplace-Dirichlet Rule-Based algorithm with *α*_epi_ = *−*60^*°*^, *α*_endo_ = 60^*°*^, *β*_epi_ = 20^*°*^and *β*_endo_ = *−*20^*°*^[42].

### 3.3 Branched Latent Neural Maps

We construct a geometry-specific surrogate model of cardiac function by building a feedforward partially-connected NN that explores the variability of our physics-based electrophysiology model detailed in Section 3.2 while structurally separating the role of temporal *t* and functional ***θ***_EP_ parameters. This recent scientific machine learning tool, proposed in [53], allows for different levels of disentanglement between inputs and outputs. The surrogate model reads:

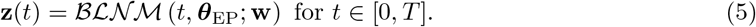

Weights and biases 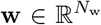 encode the algebraic structures of a feedforward partially-connected NN, which represents a map :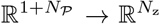from time *t* and *N*_*𝒫*_ = 7 cell-to-organ scale electrophysiology parameters 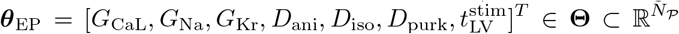 to an output vector 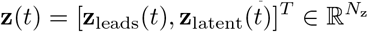 . This vector contains in silico precordial and limb leads recordings **z**_leads_(*t*) = [*V*_1_(*t*), *V*_2_(*t*), *V*_3_(*t*), *V*_4_(*t*), *V*_5_(*t*), *V*_6_(*t*), *LA*(*t*), *RA*(*t*), *F* (*t*)]^*T*^ ∈ ℝ^9^, where we use the original *LA*(*t*), *RA*(*t*) and *F* (*t*) limb leads in place of the bipolar and augmented limb leads in order to reduce the dimensionality of the output. Indeed, we reconstruct *I*(*t*), *II*(*t*), *III*(*t*), *aV L*(*t*), *aV R*(*t*) and *aV F* (*t*) a posteriori by means of Equations (3) and (4). Furthermore, vector **z**(*t*) leverages some **z**_latent_(*t*) latent variables that enhance the learned temporal dynamics by acting in regions with steep gradients [53].

We perform nonlinear optimization with the BFGS algorithm to tune NN parameters. In particular, we monitor the MSE of surrogate vs. physics-based ECG pseudopotentials to find an optimal set of weights and biases **w**, that is:

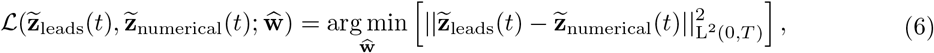

where 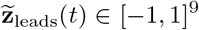 represents BLNM outputs and 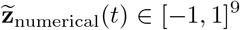 defines the physicsbased numerical simulations, both in non-dimensional form. Time 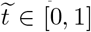 and model parameters 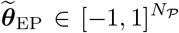 are also normalized during the training and testing phases. We refer to [53] for a detailed description of all the properties related to BLNMs that enable them to effectively learn complex physical processes.

### 3.4 Parameter estimation

We employ our trained BLNM to find a set of model parameters 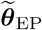 that matches **z**_ECG_(*t*) *∈* ℝ^12^ with **z**_clinical_(*t*) *∈* ℝ^12^. Here, **z**_ECG_(*t*) is the vector of BLNM physical outputs **z**_leads_(*t*) manipulated according to Equations (3) and (4) to generate the full 12-lead ECGs, and **z**_clinical_(*t*) is the clinically measured ECG data vector. In particular, we perform derivative-free optimization by employing the Nelder-Mead method [33], where we specify a loss function given by the MSE of the mismatch between the trained surrogate vs. clinical ECG potentials, that is:

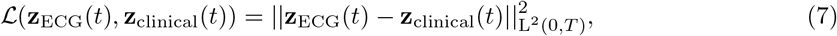

which leads to a set of calibrated model parameters 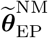 and corresponding 12-lead ECGs 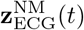. We initialize our optimization algorithm with a random set of model parameters 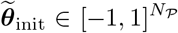 . We repeat the optimization process 100 times and we average the model parameters obtained during the different trials in order to get 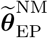.

### 3.5 Sensitivity analysis

We perform a variance-based sensitivity analysis using Shapley effects [56] in order to quantify the importance of each model parameter in fitting patient-specific 12-lead ECGs during the inference process.

Specifically, we employ Sklar’s theorem [12] to define the input multivariate distribution, which is given by a Gaussian copula and a series of *N*_*P*_ marginals defined by standard normal distributions centered in 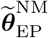, that is 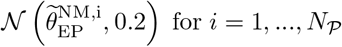 for *i* = 1, …, *N*_*𝒫*_.

Due to the high computational costs associated with testing all the different combinations of the features, we consider the random (rather than the exact) version of the algorithm to compute Shapley values. We monitor the expected marginal contribution of each model parameter to the BLNM prediction with respect to observations, that is the MSE of Equation (7). We fix 2000 permutations, 500 bootstrapped samples, and 50 samples to estimate conditional variance for 3 times.

### 3.6 Uncertainty quantification

We employ a BLNM within HMC [5] to calibrate model parameters and to perform inverse uncertainty by matching observed 12-lead ECGs from patient-specific recordings. HMC is a Markov Chain Monte Carlo (MCMC) method that aims at finding an approximation of the posterior distribution 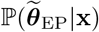, given a certain prior probability distribution 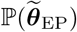 with respect to the model parameters in non-dimensional form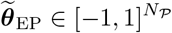. Specifically, we employ the No-U-Turn Sampler (NUTS) extension of HMC, which automatically adapts the number of steps to estimate the posterior distribution [21]. This algorithm, which shares and enhances some of the features of sequential [26] and differential evolution [61] MCMC, works well with high-dimensional target distributions, possibly presenting correlated dimensions. Moreover, HMC reaches convergence using a reduced amount of samples with respect to vanilla MCMC [21]. For further details about the mathematical derivation of HMC and its application to cardiac simulations we refer to [54].

We run 4 chains with 1,000 adaptation samples in the warm-up phase and 1,000 effective samples to estimate the posterior distribution, with a fixed 90% acceptance rate. For all model parameters, we consider prior distributions

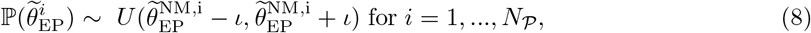

where 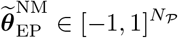is the initial guess obtained with the Nelder-Mead method. We always make sure that model parameters reside within the [ ℒ1, 1] range. We set *ι* = 0.2. Even though NUTS allows for many different initialization protocols, such as maximum a posteriori (MAP) or maximum likelihood estimation (MLE), we consider an initial random seed for each chain. This is motivated by the sensitivity of MAP and MLE over multiple runs, especially when several model parameters are calibrated with respect to noisy or highly varying time-dependent QoIs, which is the case for ECG recordings.

Several sources of uncertainty can be considered. These include model uncertainties (e.g., the discrepancy between the actual physical phenomenon and the high-fidelity model), the discretization error introduced when solving the differential equations, the surrogate modeling error of reduced-order models, and the measurement errors that intrinsically affect clinical ECG recordings (i.e., the sensitivity of the instrument used during the clinical test, variations in lead placement position by clinicians, and patient-specific factors such as breathing and motion). However, in our inverse uncertainty quantification process, we only include the measurement error and the approximation error introduced by the transition from the high-fidelity model to the BLNM-based surrogate model. In particular, we consider a multivariate normal distribution centered in the BLNM predictions **z**_ECG_(*t*) for the given patient-specific observations **z**_clinical_(*t*), which reads:

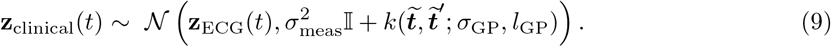

*σ*_meas_ = 0.1 is the a priori fixed standard deviation dictating the measurement error [15, 69], whereas 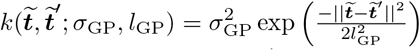 is the exponentiated quadratic kernel of a zero-mean Gaussian process 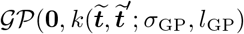 [46]. Amplitude *σ*_GP_ *∼ 𝒩*(0.01, 1.0) and correlation length *l*_GP_ *∼ 𝒩*(0.01, 1.0) are additional hyperpriors tuned during HMC to quantify the surrogate modeling error, which may change according to the specific observation. Vectors 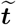 and 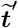 represent discrete time points in the [0, 1] interval.

A full covariance matrix in the multivariate normal distribution allows us to model the correlation among different leads. We evaluate convergence of the HMC chains by checking that the GelmanRubin diagnostic provides a value less than 1.1 for all the model parameters 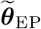, *l*_GP_ and *σ*_GP_ [6,17, 65].

### 3.7 Software and hardware

All electrophysiology simulations are performed at the Stanford Research Computing Center using `svFSIplus` [70], a C++ high-performance computing multiphysics and multiscale finite element solver for cardiac and cardiovascular modeling. This solver is part of the SimVascular software suite for patient-specific cardiovascular modeling [64].

We train the NNs by using `BLNM.jl` [22, 45], Julia library for scientific machine learning which is publicly available under MIT License at https://github.com/StanfordCBCL/BLNM.jl. This library leverages `Hyperopt.jl` [3] for parallel hyperparameter optimization by combining the Message Passing Interface (MPI) with Open Multi-Processing (OpenMP) on physical and virtual cores, respectively.

We perform sensitivity analysis and parameter estimation with uncertainty quantification using `GlobalSensitivity.jl` [11] and `Turing.jl` [16], respectively, which both exploit OpenMP and vectorized operations to speed-up computations. The code for sensitivity analysis and Bayesian parameter estimation is available within `BLNM.jl` as a test case.

Furthermore, this public repository contains the dataset encompassing all the electrophysiology simulations used for the training and testing phases, along with the patient-specific 12-lead ECGs.

## 4 Discussion

We present a complete computational pipeline to build digital twins of cardiac electrophysiology for congenital heart disease in pediatrics. This cohort of patients is understudied in cardiology [32, 62], as multiphysics and multiscale numerical simulations are mostly focused on adults with certain sets of pathologies, such as dilated, ischemic and hypertrophic cardiomyopathy, arrhythmias or bundle branch block [36, 38, 52, 58].

In this pipeline, we leverage biophysically detailed and anatomically accurate computational electrophysiology models, a recently proposed scientific machine learning tool for surrogate modeling, and robust Bayesian inference methods for personalized calibration of model parameters to match clinical 12-lead ECGs of an HLHS pediatric patient. We certify the impact and reliability of our estimation against clinical recordings by integrating fast and effective sensitivity analysis and uncertainty quantification. We run electrophysiology simulations with the estimated model parameters in order to investigate different scenarios of clinical interest in silico. We conclude that this pediatric patient presents activation and repolarization patterns similar to a left bundle branch block, where the interventricular dyssynchrony and the geometrical personalization of the Purkinje network play a minor role with respect to conductances and conductivities, even for QRS complex calibration.

Image processing allows us to get all the anatomy-specific features of this pediatric patient and our calibration of cell-to-organ level model parameters enables patient-specific electrophysiology simulations. Nevertheless, given the non-convexity of the optimization problem, it is important to stress that the final set of model parameters might not be unique and there could be other choices that lead to similar approximation errors against clinical recordings. Indeed, we notice that changing random seeds or trying different optimizers, such as second-order local BFGS or even global Adaptive Differential Evolution [68], may have an influence on the initial parameter estimation. These options are available within the `BLNM.jl` library. However, these effects are accounted for and mitigated by averaging many different trials and by running uncertainty quantification.

Performing ad-hoc sensitivity analysis for a specific parameter calibration provides individualized information, as these assessments may change on a patient to patient basis. Furthermore, we underline that sensitivity and practical identifiability (or trustworthiness) of model parameters are generally correlated. For instance, the maximum rapid delayed rectifier current conductance *G*_Kr_ and the level of interventricular dyssynchrony 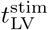 have the lowest relative impact on this 12-lead ECGs personalization (see Figure 4) and present the highest degree of uncertainty, that is a wider posterior distribution, among all physics-based model parameters (see Figure 5).

While the computational pipeline encompasses several rigorous steps, the physics-based model still requires high-performance computing and longer computational times compared to other approaches for digital twinning on adults [19, 20], which rely on more phenomenologically-based models but do not include robust methods for sensitivity analysis and uncertainty quantification. However, the monodomain equation, coupled with the ten Tusscher-Panfilov ionic model, provides an accurate mathematical model, where relevant model parameters with a direct physiological interpretation can be properly tuned. Moreover, its higher computational costs could be mitigated in future work by novel numerical methods in the framework of matrix-free [1] and Isogeometric Analysis [7].

A limitation of the presented approach lies in the lack of experimental validation of the parameter calibration process. Indeed, mathematical modeling of congenital heart disease requires several assumptions due to the current lack of information in pediatric populations regarding fiber orientation, Purkinje structure, ionic current conductances, and conduction velocities. Future studies should incorporate these data as they become available. Nevertheless, our estimations are robust, account for uncertainty quantification and are widely contained within the range explored by the electrophysiology simulations (see Table 1 and 3, along with Figure 5).

In future developments, we aim to encode anatomical variability and different CHDs, such as Tetralogy of Fallot, transposition of great arteries or atrial and ventricular septal defects within BLNMs. In this manner, the computationally expensive offline phase dictated by accurate numerical simulations and the training of the NN can be performed only once before being applied to new patients. Robust parameter estimation and uncertainty quantification will be then feasible for those CHDs within minutes, compatible with the time frame required by the clinical practice.

## Acknowledgements

We acknowledge the NSF grants 1663671 and 2105345, the NIH grants R01LM013120 and R01EB029362 and the Additional Ventures Foundation AVCC-200217. We thank Professor Francisco Sahli Costabal for the insightful discussions about Purkinje network personalization.

## Notes

https://github.com/StanfordCBCL/BLNM.jl

### Competing Interest Statement

The authors have declared no competing interest.

## References

[1] P. C. Africa, M. Salvador, P. Gervasio, L. Dede’, and A. Quarteroni. “A matrix–free high–order solver for the numerical solution of cardiac electrophysiology”. In: Journal of Computational Physics 478 (2023), p. 111984.

[2] H. Arevalo, F. Vadakkumpadan, E. Guallar, et al. “Arrhythmia risk stratification of patients after myocardial infarction using personalized heart models”. In: Nature Communications 7 (2016), pp. 113–128.

[3] F. Bagge Carlson. “Hyperopt.jl: Hyperparameter optimization in Julia”. In: (2018).

[4] D. C. Bartos, E. Grandi, and C. M. Ripplinger. “Ion Channels in the Heart”. In: Comprehensive Physiology. 2015, pp. 1423–1464.

[5] M. Betancourt and M. Girolami. “A Conceptual Introduction to Hamiltonian Monte Carlo”. In: arXiv:1701.02434 (2017).

[6] S. P. Brooks and A. Gelman. “General Methods for Monitoring Convergence of Iterative Simulations”. In: Journal of Computational and Graphical Statistics 7.4 (1998), pp. 434–455.

[7] M. Bucelli, M. Salvador, L. Dede’, and A. Quarteroni. “Multipatch Isogeometric Analysis for Electrophysiology: Simulation in a Human Heart”. In: Computer Methods in Applied Mechanics and Engineering 376 (2021), p. 113666.

[8] H. Chubb, A. Bulic, D. Mah, J. P. Moore, J. Janousek, J. Fumanelli, S. Y. Asaki, A. Pflaumer, A. C. Hill, C. Escudero, and et al. “Impact and modifiers of ventricular pacing in patients with single ventricle circulation”. In: Journal of the American College of Cardiology 80 (2022), pp. 902–914.

[9] J. Corral-Acero, F. Margara, M. Marciniak, C. Rodero, and et al. “The ‘Digital Twin’ to enable the vision of precision cardiology”. In: European Heart Journal 41.48 (2020), pp. 4556–4564.

[10] G. Del Corso, R. Verzicco, and F. Viola. “Sensitivity analysis of an electrophysiology model for the left ventricle”. In: Journal of The Royal Society Interface 17.171 (2020), p. 20200532.

[11] V. K. Dixit and C. Rackauckas. “GlobalSensitivity. jl: Performant and Parallel Global Sensitivity Analysis with Julia”. In: Journal of Open Source Software 7.76 (2022), p. 4561.

[12] F. Durante, J. Fernández-Sánchez, and C. Sempi. “A topological proof of Sklar’s theorem”. In: Applied Mathematics Letters 26.9 (2013), pp. 945–948.

[13] M. Fedele, R. Piersanti, F. Regazzoni, M. Salvador, P. C. Africa, M. Bucelli, A. Zingaro, L. Dede’, and A. Quarteroni. “A comprehensive and biophysically detailed computational model of the whole human heart electromechanics”. In: Computer Methods in Applied Mechanics and Engineering 410 (2023), p. 115983.

[14] J. A. Feinstein, D. W. Benson, A. M. Dubin, and et al. “Hypoplastic Left Heart Syndrome: Current Considerations and Expectations”. In: Journal of the American College of Cardiology 59.i1, Supplement (2012), S1–S42.

[15] F. De la Garza Salazar, M. E. Romero Ibarguengoitia, J. R. Azpiri López, and A. González Cantú. “Optimizing ECG to detect echocardiographic left ventricular hypertrophy with computerbased ECG data and machine learning”. In: PLOS ONE 16 (2021), pp. 1–14.

[16] H. Ge, K. Xu, and Z. Ghahramani. “Turing: a language for flexible probabilistic inference”. In: International Conference on Artificial Intelligence and Statistics, AISTATS 2018, 9-11 April 2018, Playa Blanca, Lanzarote, Canary Islands, Spain. 2018, p. 1682–1690.

[17] A. Gelman and D. B. Rubin. “Inference from Iterative Simulation Using Multiple Sequences”. In: Statistical Science 7 (1992), pp. 457–472.

[18] T. Gerach, S. Schuler, J. Fröhlich, et al. “Electro-Mechanical Whole-Heart Digital Twins: A Fully Coupled Multi-Physics Approach”. In: Mathematics 9.11 (2021).

[19] K. Gillette, M. Gsell, A. Prassl, E. Karabelas, U. Reiter, G. Reiter, T. Grandits, C. Payer, D. Štern, M. Urschler, J. Bayer, C. Augustin, A. Neic, T. Pock, E. Vigmond, and G. Plank. “A Framework for the generation of digital twins of cardiac electrophysiology from clinical 12-leads ECGs”. In: Medical Image Analysis 71 (2021), p. 102080.

[20] T. Grandits, J. Verhülsdonk, G. Haase, A. Effland, and S. Pezzuto. “Digital twinning of cardiac electrophysiology models from the surface ECG: a geodesic backpropagation approach”. In: arXiv:2308.08410 (2023).

[21] M. D. Homan and A. Gelman. “The No-U-Turn Sampler: Adaptively Setting Path Lengths in Hamiltonian Monte Carlo”. In: Journal of Machine Learning Research 15.1 (2014), pp. 1593–1623.

[22] M. Innes. “Flux: Elegant Machine Learning with Julia”. In: Journal of Open Source Software (2018).

[23] F. Isensee, P. F. Jaeger, S. A. A. Kohl, J. Petersen, and K. Maier-Hein. “nnU-Net: a self-configuring method for deep learning-based biomedical image segmentation”. In: Nature Methods 18 (2020), pp. 203 –211.

[24] J. Joyce, E. T. O’Leary, M. D. Y. D. M. Harrild, and J. Rhodes. “Cardiac resynchronization therapy improves the ventricular function of patients with fontan physiology”. In: American Heart Journal 230 (2020), pp. 82–92.

[25] A. Jung, M. A. F. Gsell, C. M. Augustin, and G. Plank. “An Integrated Workflow for Building Digital Twins of Cardiac Electromechanics-A Multi-Fidelity Approach for Personalising Active Mechanics”. In: Mathematics 10.5 (2022).

[26] N. Kantas, A. Doucet, S. S. Singh, and J. M. Maciejowski. “An Overview of Sequential Monte Carlo Methods for Parameter Estimation in General State-Space Models”. In: IFAC Proceedings Volumes 42 (2009), pp. 774–785.

[27] F. Kong and S. C. Shadden. “Automating Model Generation for Image-Based Cardiac Flow Simulation”. In: Journal of biomechanical engineering (2020).

[28] S. Krishnamoorthi, M. Sarkar, and W. S. Klug. “Numerical quadrature and operator splitting in finite element methods for cardiac electrophysiology”. In: International Journal for Numerical Methods in Biomedical Engineering 29 (11 2013), pp. 1243–1266.

[29] Y. Liu, S. Chen, L. J. Zühlke, G. C. Black, M. Choy, N. Li, and B. D. Keavney. “Global birth prevalence of congenital heart defects 1970–2017: updated systematic review and meta-analysis of 260 studies”. In: International Journal of Epidemiology 48 (2019), pp. 455–463.

[30] S. Longobardi, A. Lewalle, S. Coveney, I. Sjaastad, E. K. S. Espe, W. E. Louch, C. Musante, A. Sher, and S. A. Niederer. “Predicting left ventricular contractile function via Gaussian process emulation in aortic-banded rats”. In: Philosophical Transactions of the Royal Society A: Mathematical, Physical and Engineering Sciences 378.2173 (2020), p. 20190334.

[31] W. E. Lorensen and H. E. Cline. “Marching cubes: A high resolution 3D surface construction algorithm”. In: Proceedings of the 14th annual conference on computer graphics and interactive techniques (1987).

[32] A. L. Marsden and J. A. Feinstein. “Computational modeling and engineering in pediatric and congenital heart disease”. In: Current Opinion in Pediatrics 27 (2015), pp. 587–596.

[33] J. A. Nelder and R. Mead. “A Simplex Method for Function Minimization”. In: The Computer Journal 7.4 (1965), pp. 308–313.

[34] S. A. Niederer, J. Lumens, and N. A. Trayanova. “Computational models in cardiology”. In: Nature Reviews Cardiology 16 (2019), pp. 100–111.

[35] L. Pegolotti, M. R. Pfaller, N. L. Rubio, K. Ding, R. B. Brufau, E. Darve, and A. L. Marsden. “Learning Reduced-Order Models for Cardiovascular Simulations with Graph Neural Networks”. In: arXiv:2303.07310 (2023).

[36] M. Peirlinck, F. Sahli Costabal, and E. Kuhl. “Sex differences in drug-induced arrhythmogenesis”. In: Frontiers in Physiology 12 (2021), p. 708435.

[37] M. Peirlinck, F. Sahli Costabal, K. Sack, J. Choy, G. Kassab, J. Guccione, M. De Beule, P. Segers, and E. Kuhl. “Using machine learning to characterize heart failure across the scales.” In: Biomechanics and Modeling in Mechanobiology 18 (2019), 1987–2001.

[38] M. Peirlinck, F. Sahli Costabal, J. Yao, J. Guccione, S. Tripathy, Y. Wang, D. Ozturk, P. Segars, T. Morrison, S. Levine, and E. Kuhl. “Precision medicine in human heart modeling”. In: Biomechanics and Modeling in Mechanobiology 20 (2021), pp. 803–831.

[39] M. Peirlinck, J. Yao, F. Sahli Costabal, and E. Kuhl. “How drugs modulate the performance of the human heart”. In: Computational Mechanics 69 (2022), pp. 1397–1411.

[40] P. Pentenga, A. Stroh, W. van Genuchten, W. Helbing, and M. Peirlinck. “Shape Morphing and Slice Shift Correction in Congenital Heart Defect Model Generation”. In: International Conference on Functional Imaging and Modeling of the Heart (2023), pp. 347–355.

[41] M. Pfaller, J. Hörmann, M. Weigl, et al. “The importance of the pericardium for cardiac biomechanics: from physiology to computational modeling”. In: Biomechanics and Modeling in Mechanobiology 18 (2019), pp. 503–529.

[42] R. Piersanti, P. C. Africa, M. Fedele, et al. “Modeling cardiac muscle fibers in ventricular and atrial electrophysiology simulations”. In: Computer Methods in Applied Mechanics and Engineering 373 (2021), p. 113468.

[43] A. Prakosa, H. Arevalo, D. Dongdong, et al. “Personalized virtual-heart technology for guiding the ablation of infarct-related ventricular tachycardia”. In: Nature Biomedical Engineering 2 (2018), pp. 732–740.

[44] A. Quarteroni, L. Dede’, A. Manzoni, and C. Vergara. Mathematical Modelling of the Human Cardiovascular System: Data, Numerical Approximation, Clinical Applications. Cambridge University Press, 2019.

[45] C. Rackauckas and Q. Nie. “Differentialequations.jl–a performant and feature-rich ecosystem for solving differential equations in julia”. In: Journal of Open Research Software 5.1 (2017), p. 15.

[46] C. E. Rasmussen and C. K. I. Williams. Gaussian Processes for Machine Learning. The MIT Press, 2005.

[47] F. Regazzoni, M. Salvador, L. Dede’, and A. Quarteroni. “A machine learning method for realtime numerical simulations of cardiac electromechanics”. In: Computer Methods in Applied Mechanics and Engineering 393 (2022), p. 114825.

[48] F. Sahli Costabal, D. E. Hurtado, and E. Kuhl. “Generating Purkinje networks in the human heart”. In: Journal of biomechanics 49 (2016), pp. 2455–2465.

[49] F. Sahli Costabal, K. Matsuno, J. Yao, P. Perdikaris, and E. Kuhl. “Machine learning in drug development: Characterizing the effect of 30 drugs on the QT interval using Gaussian process regression, sensitivity analysis, and uncertainty quantification”. In: Computer Methods in Applied Mechanics and Engineering 348 (2019), pp. 313–333.

[50] F. Sahli Costabal, J. Yao, and E. Kuhl. “Predicting the cardiac toxicity of drugs using a novel multiscale exposure–response simulator”. In: Computer methods in biomechanics and biomedical engineering 21 (2018), pp. 232–246.

[51] F. Sahli Costabal, J. Yao, and E. Kuhl. “Predicting drug-induced arrhythmias by multiscale modeling”. In: International Journal for Numerical Methods in Biomedical Engineering 34.5 (2018), e2964.

[52] M. Salvador, M. Fedele, P. C. Africa, et al. “Electromechanical modeling of human ventricles with ischemic cardiomyopathy: numerical simulations in sinus rhythm and under arrhythmia”. In: Computers in Biology and Medicine 136 (2021), p. 104674.

[53] M. Salvador and A. L. Marsden. “Branched Latent Neural Maps”. In: Computer Methods in Applied Mechanics and Engineering 418 (2024), p. 116499.

[54] M. Salvador, F. Regazzoni, L. Dede’, and A. Quarteroni. “Fast and robust parameter estimation with uncertainty quantification for the cardiac function”. In: Computer Methods and Programs in Biomedicine 231 (2023), p. 107402.

[55] H. Si. “TetGen, a Delaunay-Based Quality Tetrahedral Mesh Generator”. In: ACM Transactions on Mathematical Software (TOMS) 41 (2015), pp. 1 –36.

[56] E. Song, B. L. Nelson, and J. Staum. “Shapley Effects for Global Sensitivity Analysis: Theory and Computation”. In: SIAM/ASA Journal on Uncertainty Quantification 4.1 (2016), pp. 1060–1083.

[57] M. Strocchi, C. M. Augustin, M. A. F. Gsell, et al. “A publicly available virtual cohort of four-chamber heart meshes for cardiac electro-mechanics simulations”. In: PLOS ONE 15 (2020), pp. 1–26.

[58] M. Strocchi, K. Gillette, A. Neic, M. K. Elliott, N. Wijesuriya, V. Mehta, E. J. Vigmond, G. Plank, C. A. Rinaldi, and S. A. Niederer. “Comparison between conduction system pacing and cardiac resynchronization therapy in right bundle branch block patients”. In: Frontiers in Physiology 13 (2022), p. 1011566.

[59] M. Strocchi, S. Longobardi, C. M. Augustin, M. A. F. Gsell, A. Petras, C. A. Rinaldi, E. J. Vigmond, G. Plank, C. J. Oates, R. D. Wilkinson, and S. A. Niederer. “Cell to Whole Organ Global Sensitivity Analysis on a Four-chamber Electromechanics Model Using Gaussian Processes Emulators”. Submitted to PLOS Computational Biology.

[60] C. Sánchez, G. D’Ambrosio, F. Maffessanti, E. G. Caiani, F. W. Prinzen, R. Krause, A. Auricchio, and M. Potse. “Sensitivity analysis of ventricular activation and electrocardiogram in tailored models of heart-failure patients”. In: Medical & Biological Engineering & Computing 56.3 (2018), pp. 491–504.

[61] C. J. F. Ter Braak. “A Markov Chain Monte Carlo version of the genetic algorithm Differential Evolution: easy Bayesian computing for real parameter spaces”. In: Statistics and Computing 16 (2006), pp. 239–249.

[62] O. Z. Tikenogullari, M. Peirlinck, H. Chubb, A. M. Dubin, E. Kuhl, and A. L. Marsden. “Effects of cardiac growth on electrical dyssynchrony in the single ventricle patient”. In: Computer Methods in Biomechanics and Biomedical Engineering 0.0 (2023), pp. 1–17.

[63] K. H. ten Tusscher and A. V. Panfilov. “Alternans and spiral breakup in a human ventricular tissue model”. In: American Journal of Physiology. Heart and Circulatory Physiology 291 (2006), pp. 1088–1100.

[64] A. Updegrove, N. M. Wilson, J. Merkow, H. Lan, A. L. Marsden, and S. C. Shadden. “Sim-Vascular: An Open Source Pipeline for Cardiovascular Simulation”. In: Annals of Biomedical Engineering 45 (2017), 525–541.

[65] D. Vats and C. Knudson. “Revisiting the Gelman-Rubin Diagnostic”. In: arXiv:1812.09384 (2018).

[66] B. Winkler, C. Nagel, N. Farchmin, S. Heidenreich, A. Loewe, O. Dössel, and M. Bär. “Global Sensitivity Analysis and Uncertainty Quantification for Simulated Atrial Electrocardiograms”. In: Metrology 3.1 (2022), 1–28.

[67] X. Xu, T. Wang, Z. Jian, H. Yuan, M. Huang, J. Cen, Q. Jia, Y. Dong, and Y. Shi. “ImageCHD: A 3D Computed Tomography Image Dataset for Classification of Congenital Heart Disease”. In: International Conference on Medical Image Computing and Computer-Assisted Intervention. 2020.

[68] J. Zhang and A. C. Sanderson. “JADE: Adaptive Differential Evolution With Optional External Archive”. In: IEEE Transactions on Evolutionary Computation 13.5 (2009), pp. 945–958.

[69] S. H. Zhou, E. D. Helfenbein, J. M. Lindauer, R. E. Gregg, and D. Q. Feild. “Philips QT interval measurement algorithms for diagnostic, ambulatory, and patient monitoring ECG applications”. In: Annals of Noninvasive Electrocardiology 14(Suppl 1) (2009), S3–S8.

[70] C. Zhu, V. Vedula, D. Parker, N. Wilson, S. Shadden, and A. Marsden. “svFSI: A Multiphysics Package for Integrated Cardiac Modeling”. In: Journal of Open Source Software 7.78 (2022), p. 4118.

